# A highly divergent mitochondrial genome in extant Cape buffalo from Addo Elephant National Park, South Africa

**DOI:** 10.1101/2023.01.14.524049

**Authors:** Deon de Jager, Marlo Möller, Eileen Hoal, Paul van Helden, Brigitte Glanzmann, Cindy Harper, Paulette Bloomer

## Abstract

The reduced cost of next-generation sequencing (NGS) has allowed researchers to generate nuclear and mitochondrial genome data to gain deeper insights into the phylogeography, evolutionary history, and biology of non-model species. While the Cape buffalo (*Syncerus caffer caffer*) has been well-studied across its range with traditional genetic markers over the last 25 years, researchers are building on this knowledge by generating whole genome, population-level data sets to improve understanding of the genetic composition and evolutionary history of the species. Using publicly available NGS data, we assembled 40 Cape buffalo mitochondrial genomes (mitogenomes) from four protected areas in South Africa, expanding the geographical range and almost doubling the number of mitogenomes available for this species. Coverage of the mitogenomes ranged from 154-1,036X. Haplotype and nucleotide diversity for Kruger National Park (n = 15) and Mokala National Park (n = 5) were similar to diversity levels in southern and eastern Africa. Hluhluwe-iMfolozi Park (n = 15) had low levels of genetic diversity, with only four haplotypes detected, reflecting its past bottleneck. Addo Elephant National Park (n = 5) had the highest nucleotide diversity of all populations across Africa, which was unexpected, as it is known from previous studies to have low nuclear diversity. This diversity was driven by a highly divergent mitogenome from one sample, which was subsequently identified in another sample via Sanger sequencing of the cytochrome *b* gene. Using a fossil-calibrated phylogenetic analysis, we estimated that this lineage diverged from all other Cape buffalo lineages approximately 2.51 million years ago. We discuss several potential sources of this mitogenome but propose that it most likely originated through introgressive hybridisation with an extinct buffalo species, either *S. acoelotus* or *S. antiquus*. We conclude by discussing the conservation consequences of this finding for the Addo Elephant National Park population, proposing careful genetic management to prevent inbreeding depression while maintaining this highly unique diversity.

## Introduction

The Cape buffalo (*Syncerus caffer caffer*, Sparrman 1779) is a charismatic African bovid known for being one of the “Big Five”. It is also a reservoir for important veterinary diseases, such as foot-and-mouth disease, Corridor disease, brucellosis, and bovine tuberculosis (Laubscher & Hoffman, 2012), and is a high-value species in both hunting and wildlife ranching, particularly in South Africa (Taylor, Lindsey, & Davies-Mostert, 2016). Consequently, this species has been well-studied from ecological, disease, and genetic perspectives throughout most of its range (Cornélis et al., 2014; de Jager et al., 2021; de Jager, Harper, & Bloomer, 2020; Heller, Lorenzen, Okello, Masembe, & Siegismund, 2008; Heller, Okello, & Siegismund, 2010; O’Ryan et al., 1998; Quinn et al., 2023; Simonsen, Siegismund, & Arctander, 1998; Smitz et al., 2013; Smitz et al., 2014; van Hooft, Groen, & Prins, 2000, 2002, 2003).

However, most of the genetic studies have been based on microsatellite loci, which are difficult to combine and compare between studies, or on a portion of the control region (CR) of the mitochondrial genome (mitogenome), which is easier to combine across studies, but is usually a short segment that might not be representative of the rest of the mitogenome. With the ever-decreasing cost of next-generation sequencing (NGS), it is possible to generate population-level whole genome sequences for non-model species, as illustrated for Cape buffalo in South Africa by de Jager et al. (2021), and across much of the rest of the range of the subspecies by Quinn et al., (2023). In their study, Heller, Brüniche-Olsen, and Siegismund (2012) generated population-level mitogenomes from populations in Ethiopia, Kenya, Zimbabwe, and Botswana. These types of data are easier to combine across studies and are more powerful than traditional nuclear and mitochondrial markers. However, some gaps in genomic data still exist, for example the lack of nuclear genomes from the other African buffalo subspecies, and the lack of Cape buffalo mitogenomes from South Africa.

In the latest Red List assessment by the International Union for Conservation of Nature (IUCN), the status of the African buffalo (*S. caffer*) changed from Least Concern to Near Threatened, predominantly due to ongoing and predicted population declines (IUCN, 2019). However, Cape buffalo is regionally classified as Least Concern in southern Africa and is increasing in numbers in South Africa (Tambling, Venter, du Toit, & Child, 2016). In this study, we present 40 Cape buffalo mitogenomes from four South African protected areas to fill the gap in mitogenomic data for the species. We analyse these in the context of previously published mitogenomes to compare mitochondrial genome diversity across a large part of the Cape buffalo distribution range and discuss the conservation implications of our findings. With the data presented here (and in previous studies) we aim to ultimately assist in the genetic management and conservation of Cape buffalo populations within, and beyond, South Africa. This is particularly relevant given the importance of genetic assessment and monitoring of wildlife populations now recognised at national and international levels (Hoban et al., 2020; Hoban et al., 2021; Laikre et al., 2020).

## Methods

### Mitogenome assembly and annotation

Mitogenomes were assembled using NOVOPlasty v4.0 (Dierckxsens, Mardulyn, & Smits, 2017) from the cleaned and trimmed reads of 40 Cape buffalo nuclear genomes sequenced by de Jager et al. (2021). NOVOPlasty *de novo* assembles organellar genomes from whole genome sequencing data using a seed-and-extend algorithm (Dierckxsens et al., 2017). The 40 buffalo samples originate from four protected areas in South Africa, namely Addo Elephant National Park (NP) (n = 5), Hluhluwe-iMfolozi Park (n = 15), Kruger NP (n = 15), and Mokala NP (n = 5); the latter is an introduced population established with Kruger buffalo (de Jager et al., 2020). We used the mitogenome of a Cape buffalo from Masai Mara National Reserve in Kenya (GenBank accession: JQ235542 (Heller et al., 2012)) as the seed for the NOVOPlasty assembly. Genome range was set to 15,000-18,000 base pairs (bp) and k-mer set to 33. NOVOPlasty parameter settings were the same for all samples and are provided in a configuration file, an example of which is available here: https://github.com/DeondeJager/Buffalo_Mitogenomics. The assembled mitogenomes were manually checked for ambiguous bases in Geneious Prime v2021.2.2. For the 12 mitogenomes that had one or more ambiguous bases, the cleaned and trimmed genome reads were mapped against the assembled mitogenome using the BWA-MEM algorithm in bwa v0.1.17 (Li & Durbin, 2009). The resultant BAM file was sorted by position using SAMtools v1.9 (Li et al., 2009) and subsequently viewed in Integrative Genomics Viewer (IGV) v2.5.3 (Robinson et al., 2011; Thorvaldsdóttir, Robinson, & Mesirov, 2012) and ambiguities manually resolved where possible. The assembled mitogenomes were *de novo* annotated with MITOS2 (Donath et al., 2019). The MITOS2 annotations were manually curated through comparisons with annotations of the cattle (*Bos taurus*) RefSeq reference mitogenome (NC_006853.1) and the African buffalo reference mitogenome (NC_020617.1) (Hassanin et al. 2012). Specifically, all protein-coding genes were translated in Geneious using the vertebrate mitochondrial codon table (transl_table 2) and the protein lengths were compared to those from the two reference mitogenomes. The mitogenome nucleotide sequence data reported are available in the Third Party Annotation Section of the DDBJ/ENA/GenBank databases under the accession numbers TPA: BK062533-BK062572.

### Genetic diversity and structure

To investigate whether there was any sub-structuring of mitogenomes across the range of Cape buffalo, the 40 South African mitogenomes were combined with the 43 from Heller et al. (2012), originating from Ethiopia, Kenya, Zimbabwe, and Botswana, and the mitogenome from Hassanin et al. (2012) from Tanzania, totalling 84 mitogenomes (Table S1). The 13 protein-coding genes, two rRNA genes and the control region were extracted, concatenated, and aligned using MUSCLE (Edgar, 2004). The alignments were imported into MEGA X (Kumar, Stecher, Li, Knyaz, & Tamura, 2018) in FASTA format and exported in nexus format, excluding ambiguous sites, for a total alignment length of 13,786 bp. The alignment was used to construct a minimum spanning network (Bandelt, Forster, & Röhl, 1999) of haplotypes in PopArt v1.7 (http://popart.otago.ac.nz) (Leigh & Bryant, 2015). PopArt automatically masks sites with >5% missing data (gaps or ambiguous nucleotides). Diversity and divergence statistics were calculated in DnaSP v6.12.03 (Rozas et al., 2017), ignoring gaps (see Table 1 for number of sites).

**Table 1.** Summary statistics of mitogenome alignments of various subsets of the data calculated in DnaSP v6.12.03.

| **Dataset** | **Sample size (N)** | **Segregating sites (S)** | **No. of haplotypes (h)** | **Haplotype diversity (Hd)** | **Nucleotide diversity ($\pi$)** | **Average no. of nucleotide differences (k)** | **Tajima's *D*** | **No. of sites in alignment (bp)** |
| --- | --- | --- | --- | --- | --- | --- | --- | --- |
| Addo Elephant National Park | 5 | 518 | 3 | 0.8 | 0.01526 | 209.2 | -1.21246 | 13,706 |
| Addo Elephant National Park (excl. divergent) | 4 | 10 | 2 | 0.667 | 0.00049 | 6.667 | 2.22343 | 13,710 |
| Kruger National Park | 15 | 228 | 12 | 0.971 | 0.00474 | 64.971 | -0.34306 | 13,707 |
| Mokala National Park | 5 | 152 | 5 | 1 | 0.00481 | 65.9 | -0.78218 | 13,707 |
| Hluhluwe-iMfolozi Park | 15 | 95 | 4 | 0.467 | 0.00258 | 35.41 | 0.92808 | 13,705 |
| **South Africa** | **40** | **702** | **23** | **0.919** | **0.00626** | **85.813** | **-1.81535\*** | **13,702** |
| **South Africa (excl. divergent)** | **39** | **287** | **22** | **0.915** | **0.00465** | **63.682** | **-0.24381** | **13,704** |
| Botswana (Chobe NP) | 8 | 203 | 7 | 0.964 | 0.00537 | 73.607 | -0.32758 | 13,706 |
| Zimbabwe (Hwange NP) | 8 | 149 | 5 | 0.857 | 0.00451 | 61.786 | 0.37156 | 13,705 |
| Kenya (Masai Mara NR) | 20 | 228 | 14 | 0.953 | 0.00372 | 50.984 | -0.88809 | 13,704 |
| Ethiopia (Omo Valley) | 7 | 134 | 7 | 1 | 0.00342 | 46.667 | -0.89648 | 13,633 |
| **Heller et al. 2012 (recalculation)** | **43** | **389** | **30** | **0.978** | **0.00439** | **59.886** | **-1.25622** | **13,627** |
| **Heller et al. 2012 (original)** | **43** | **456** | **30** | **-** | **0.00435** | **-** | **-1.24** |  |

| Dataset | Sample size (N) | Segregating sites (S) | No. of haplotypes (h) | Haplotype diversity (Hd) | Nucleotide diversity ($\pi$) | Average no. of nucleotide differences (k) | Tajima's *D* | No. of sites in alignment (bp) |
| --- | --- | --- | --- | --- | --- | --- | --- | --- |
| Southern Africa (South Africa, Botswana, Zimbabwe) | 56 | 780 | 33 | 0.953 | 0.00604 | 82.764 | -1.85935* | 13,699 |
| Southern Africa (excl. divergent) | 55 | 382 | 32 | 0.951 | 0.0049 | 67.141 | -0.71192 | 13,701 |
| East Africa (Tanzania, Kenya, Ethiopia) | 28 | 277 | 21 | 0.974 | 0.00362 | 49.323 | -1.23484 | 13,629 |
| All mitogenomes | 84 | 857 | 53 | 0.976 | 0.00517 | 70.476 | -2.04442* | 13,624 |
| All mitogenomes (excl. divergent) | 83 | 488 | 52 | 0.975 | 0.00444 | 60.51 | -1.32907 | 13,625 |

\* $P < 0.05$

The control region (CR) of the 84 Cape buffalo mitogenomes were extracted and combined with publicly available CR sequences on GenBank of all African buffalo (*S. caffer*) subspecies from Simonsen et al. (1998), van Hooft et al. (2002), and Smitz et al. (2013), giving a final dataset of 876 sequences (Table S2). Sequences were aligned using MAFFT (Katoh & Standley, 2013) and cleaned and trimmed using Gblocks (Castresana, 2000; Talavera & Castresana, 2007), both implemented on the NGPhylogeny.fr platform (Lemoine et al., 2019). Gblocks parameters were set as follows: Minimum number of sequences for a conserved position (b1) = 50% of sequences + 1, Minimum number of sequences for a flank position (b2) = 85% of sequences, Maximum number of contiguous non-conserved positions (b3) = 8, Minimum length of a block (b4) = 10, Allowed gap positions (b5) = “With half”. The resultant alignment, consisting of 285 aligned bases, was used to construct a minimum spanning network of haplotypes in PopArt.

A Bayesian phylogenetic tree was constructed by aligning whole mitogenomes using MAFFT, with the cattle (*Bos taurus*, NC_006853.1) and water buffalo (*Bubalus bubalis*, NC_049568.1) reference mitogenomes used as outgroups, giving an alignment of 16,436 bp. The best substitution model was determined to be HKY+G+I, using the Bayesian information criterion in jModelTest v2.1.10 (Darriba, Taboada, Doallo, & Posada, 2012; Guindon & Gascuel, 2003), with 5 substitution schemes. The tree was constructed in MrBayes v3.2.7 (Ronquist et al. 2012), with two independent runs with four chains and five million generations each, sampling trees every 100 generations, and using a burn-in of 25%. Convergence was reached both within and between the two runs, as indicated by the ESS values >100, the potential scale reduction factor (PSRF) being very close to 1 for all parameters, and the standard deviation of split frequencies being less than the suggested 0.01. The consensus tree across the two runs was visualised in FigTree v1.4.4 (http://tree.bio.ed.ac.uk/software/figtree/) and further processed in InkScape v1.3.2 (https://inkscape.org/).

### Validation of the divergent mitogenome

The divergent mitogenome in sample A_268_14, a male buffalo from Addo Elephant NP, was apparent from the mitogenomic alignment and haplotype network. To ensure that this mitogenome was not an artefact due to an assembly error, or a nuclear copy of the mitogenome (NuMT), we performed several quality checks. First, it was confirmed that all the protein-coding genes were predicted to encode full-length proteins. Second, the genomic reads were mapped to the assembled mitogenome using the BWA-MEM algorithm in bwa v0.1.17 and the BAM file visualised in IGV to check for any obvious assembly errors, of which none were identified.

Finally, a segment of the cytochrome *b* gene was sequenced using Sanger sequencing, not only of the sample in question, but also of its two closest relatives from Addo Elephant NP: A_05_08 (male) and A_247_14 (female), as determined with 11 microsatellite loci in de Jager, Swarts, Harper, and Bloomer (2017). DNA was extracted from whole blood using the Qiagen DNeasy Blood & Tissue Kit (Qiagen, Hilden, Germany), according to the manufacturer’s instructions. A 354 bp internal portion of the cytochrome *b* gene was amplified using the primers L14841 and H15149 (Kocher et al., 1989) (L14841: 5’-[AAAAAGCTTCCAT]CCAACATCTCAGCATGATGAAA-3’, H15149: 5’-[AAACTGCAG]CCCCTCAGAATGATATTTGTCCTCA-3’, where the square brackets indicate extra sequence present in the primers designed by Kocher et al. (1989), but not included in the primers used in this study). PCR amplification was performed in reactions containing 1X Standard Reaction Buffer, 2.5 mM MgCl_2_, 2 mM dNTP mix, 1X bovine serum albumin, 3 U Super-Therm *Taq* DNA polymerase (Thermo Fisher Scientific, Massachusetts, United States), 0.1 μM L14841 forward primer, 0.1 μM H15149 reverse primer, 10-30 ng DNA, and molecular grade water to a final reaction volume of 25 μL, with the following conditions in a SimpliAmp Thermal Cycler (Life Technologies, California, United States): 95°C for 3 min, followed by 25 cycles of 94°C for 30 s, 55°C for 30 s, 72°C for 30 s, and a final extension step at 72°C for 7 min. PCR products were cleaned via ethanol precipitation and 2.5 µL clean product used in a cycle sequencing reaction (BigDye© Terminator v3.1 Cycle Sequencing Kit (Thermo Fisher Scientific)) containing 1X Ready Reaction Premix, 1X BigDye© Sequencing Buffer, 0.5 μM L14841 forward primer or 0.1 μM H15149 primer and molecular grade water to a final volume of 10 μL. Cycle sequencing reactions were performed in a SimpliAmp Thermal Cycler with 25 cycles at 96°C for 10 s, 50°C for 5 s and 60°C for 4 min. Products were cleaned via ethanol precipitation and submitted to the DNA Sequencing Facility of the Faculty of Natural and Agricultural Sciences at the University of Pretoria for sequencing on a 3500xl automated sequencer (Life Technologies, California, United States). All products were sequenced in both the forward and reverse directions to obtain 2X depth of coverage of each base position. The sequences were cleaned and trimmed in Geneious and the consensus sequences were aligned using MUSCLE to the cytochrome *b* genes extracted from the five assembled mitogenomes from Addo Elephant NP. The alignments were trimmed to a final length of 237 bp and used to construct a minimum spanning network of haplotypes in PopArt. The nucleotide sequences generated via Sanger sequencing are available on GenBank under the following accession numbers: OP485291-OP485293.

### Divergence dating

To estimate the time of divergence between the divergent mitogenome and other Cape buffalo mitogenomes, and to determine its phylogenetic position within Bovini, we constructed a dated phylogeny in BEAST v2.5.2 (Bouckaert et al., 2019). The 13 protein-coding genes, two rRNA genes (12S and 16S), and the CR were extracted, concatenated, and aligned using MAFFT, from the following mitogenomes: the divergent *S. c. caffer* mitogenome from Addo Elephant NP (A_268_14), another *S. c. caffer* from Addo Elephant NP (A_87_13), and a third *S. c. caffer* from Tanzania (GenBank accession: EF536353)), eight other Bovidae mitogenomes included in Hassanin et al. (2012), namely *Tragelaphus oryx* (JN632704), *Bos taurus taurus* (EU177832), *Bos taurus indicus* (EU177868), *Bos grunniens* (NC_006380), *Bison bison* (JN632601), *Bubalus bubalis bubalis* (AF547270), *Bubalus bubalis carabanesis* (JN632607), and the dwarf musk deer (*Moschus berezovskii*, NC_012694) belonging to the Moschidae family, also included in Hassanin et al. (2012), which was used as an outgroup. The alignment was defined as three partitions: the 13 protein coding genes (11,394 sites), the two rRNA genes (2,551 sites), and CR (1,027 sites). The best-fit substitution model for each partition was estimated in jModeltest v2.1.10 and determined to be GTR+G for the protein-coding and rRNA partitions, and HKY+G (Hasegawa, Kishino, & Yano, 1985) for the CR partition, all with four rate categories, based on the Bayesian information criterion.

The evolutionary model was prepared in BEAUTi v2.6.5 (Bouckaert et al., 2019), with linked clock and tree models, empirical base frequencies, a relaxed log normal clock (Drummond, Ho, Phillips, & Rambaut, 2006) with a clock rate of 1.0, and the fossilized birth-death tree model (Heath et al. 2014). Two fossil calibration points were used following Bibi (2013): Crown Bovidae, which contained all taxa except *M*. *berezovskii* and was set as a normal distribution with a mean of 18 million years ago (Mya) and a sigma of 1.0, giving a distribution with a 2.5% quantile of 16 Mya and a 97.5% quantile of 20 Mya, and Crown Bovini, which contained all taxa except *M*. *berezovskii* and *T*. *oryx* and was set as a normal distribution with a mean of 9 Mya and a sigma of 1.0, giving a distribution with a 2.5% quantile of 7.04 Mya and a 97.5% quantile of 11.0 Mya. Monophyly was enforced in both cases. The analysis was executed in two independent runs each with an MCMC chain length of 20 million generations, sampling every 1,000 runs. The xml file (available at https://github.com/DeondeJager/Buffalo_Mitogenomics) was executed in BEAST and the resulting log files were examined in Tracer v1.7.2 (Rambaut, Drummond, Xie, Baele & Suchard, 2018), which showed that all parameters converged, with individual and combined effective sample sizes greater than 200, and in most cases greater than 700. The tree files from both runs were combined using LogCombiner (Bouckaert et al., 2019) and the output was used in Tree Annotator (Bouckaert et al., 2019) to build a maximum clade credibility (MCC) tree with a 10% burn-in (4,000 trees), a posterior probability limit of 0.5, and using median node heights. The MCC tree was visualised in FigTree and further processed in InkScape.

## Results and discussion

### South African mitogenomes

The 40 South African (SA) Cape buffalo mitogenomes were each assembled into a single, circularised contig using NOVOPlasty. Coverage ranged from 154X to 1,036X (mean: 419X) and lengths were 16,357-16,362 bp (Table S3), which was similar to the sizes of published mitogenomes for the subspecies (16,357-16,361 bp) (Hassanin et al., 2012; Heller et al., 2012). The cause of these slight size differences was usually one or two base pair indels in the rRNA genes or CR. There were two exceptions to this, with a mitogenome from Kruger NP (B98_597) having two insertions of 5 bp and 3 bp in the CR, increasing its size to 16,367 bp, and a mitogenome from Addo Elephant NP (A_268_14) having a 10 bp insertion in the CR increasing its size to 16,375 bp. A large deletion of 74 bp in the CR of a Cape buffalo mitogenome from Ethiopia was previously reported by Heller et al. (2012), showing that large indels can be present in the D-loop within the subspecies.

The MITOS2 annotations were mostly accurate, as the protein-coding genes of all 40 assembled SA mitogenomes had no internal stop codons and were predicted to produce proteins of the expected length, as compared to the cattle and African buffalo reference mitogenomes. The exceptions were the genes *cox3* and *nad5*, which were predicted to produce proteins that were respectively one and three amino acids longer than the references in all 40 samples. This was due to the 3’ end of *cox3* predicted by MITOS2 to be four base pairs downstream, and the start codon of *nad5* predicted to be nine base pairs upstream, compared to the references. These annotations were manually edited to match the annotations of the reference mitogenomes. Two samples from Addo Elephant NP, A_243_14 and A_251_14, had an A to G transition in the second position of the stop codon of the *atp8* gene (*m.8326A>G*). This change was supported by 500/536 reads (93%) at this position in sample A_243_14 containing a G, with only 35 reads (7%) containing an A and one read (0%) containing a C. For sample A_251_14, 347/376 reads (92%) contained a G at this position, with only 27 reads (7%) containing an A and two reads (1%) containing a C. The substitution changes the stop codon (UAA) to one coding for the amino acid tryptophan (UGA). However, there is an in-frame stop codon six base pairs downstream (UAG), resulting in the predicted protein product being 68 amino acids long, as opposed to the expected 66 amino acids, with a tryptophan (UGA) and serine (UCC) added to the C-terminal of the ATP8 protein. The above results indicated that the mitogenomes assembled from the whole genome sequencing reads were of high quality and accuracy.

### Genetic diversity and structure

The Addo Elephant NP population had the highest nucleotide diversity of all Cape buffalo populations in this study, almost three times that of the next most diverse population (Chobe NP) (Table 1). This was surprising, as the population from Addo Elephant NP is known to have the lowest nuclear genetic diversity of all natural Cape buffalo populations studied thus far, due to a strong historical bottleneck and subsequent isolation (de Jager et al., 2021; de Jager et al., 2020; O’Ryan et al., 1998; Quinn et al., 2023). This high diversity was driven by one sample, A_268_14, the mitogenome of which was highly divergent, not only from the other Addo Elephant NP mitogenomes, but also those of the rest of the subspecies. Interestingly, the nuclear genome of this individual did not show any unique signals in analyses conducted by de Jager et al. (2021) or Quinn et al. (2023) and clustered with the nuclear genomes of the other Addo Elephant NP buffalo. The average number of nucleotide substitutions between this mitogenome and the rest of Cape buffalo (n = 83) was 0.03517, a divergence of 3.5%. When this sample was excluded, the Addo Elephant NP population had the lowest nucleotide diversity, five times lower than that of the next lowest population (Hluhluwe-iMfolozi Park) (Table 1), which is more in line with the nuclear diversity results from previous studies and the population’s demographic history. Hluhluwe-iMfolozi Park, having also experienced an historical bottleneck, though seemingly not as extreme as that of Addo Elephant NP (de Jager et al., 2021; O’Ryan et al., 1998; Quinn et al., 2023), harbours the second largest free-ranging buffalo population in South Africa (∼4,544 individuals in 2019 (EKZNW, 2019)), but had the lowest haplotype diversity with only four haplotypes from 15 samples (Table 1). Additionally, its nucleotide diversity was second lowest and substantially lower than that of Mokala NP and Kruger NP, the latter of which harbours the largest free-ranging buffalo population in South Africa (∼40,900 individuals in 2011 (Cornélis et al., 2014)). The Tajima’s *D* values of both Addo Elephant NP (excluding the divergent mitogenome) and Hluhluwe-iMfolozi Park were positive, which is potentially an indication of their recent bottlenecks, though these values were not significantly different from zero (Table 1).

Nucleotide diversity was higher in southern Africa (Zimbabwe, Botswana, SA) than in East Africa (Kenya, Ethiopia), even when excluding the divergent mitogenome from Addo Elephant NP, whereas higher haplotype diversity was observed in East Africa (Table 1). It is hypothesised, based on genetic and fossil evidence, that Cape buffalo dispersed to southern Africa from East Africa during the last ∼80,000 years (Heller et al., 2012; Smitz et al., 2013). While our nucleotide diversity results may not seem to support this scenario (where diversity is expected to be higher in the source compared to the sink population), we note that there is only one mitogenome available from Tanzania and none from Uganda. The former has the largest free-ranging Cape buffalo population, upwards of 100,000 individuals (Cornélis et al., 2014), while the region of present-day Uganda appears to harbour several ancestral mitochondrial lineages and unique haplotypes of Cape buffalo (Smitz et al., 2013). Inclusion of mitogenomes from these two locations would more accurately represent the genetic diversity of East African buffalo and likely reflect the expected source-sink pattern. The Tajima’s *D* value for southern Africa was negative and significantly different from zero (Table 1), which could be indicative of a population expansion, but when the divergent mitogenome was excluded, the value no longer deviated significantly from zero, indicating that this is what was driving this signal. The same was observed for Addo Elephant NP and South Africa (Table 1). Genetic diversity and migration rates are high throughout most of the range of the subspecies (Simonsen et al., 1998; Smitz et al., 2013; van Hooft et al., 2000), making other evolutionary patterns more difficult to discern, while recent demographic events (e.g. bottlenecks in the last ∼300 years, exacerbated by subsequent isolation) of individual populations may mask signals of previous evolutionary or demographic events (Excoffier & Schneider, 1999; Rasmus Heller et al., 2012).

The minimum spanning network of mitogenomes showed no clear evidence of haplotypes clustering according to location and reflected the high diversity within the subspecies, as indicated by few shared haplotypes and many mutational steps between haplotypes (Figure 1A). There was a tendency for haplotypes from southern Africa to be at the tips of the network, with those from East Africa generally more internal (Figure 1A), which adds support to the east-south dispersal hypothesis (Smitz et al., 2013). The Bayesian phylogenetic tree also showed no clear relationship between mitogenomes and geographical origin at a broad scale (i.e., between eastern and southern Africa), which was well supported with most nodes having a posterior probability close or equal to one (Figure S1).

**Figure 1.**
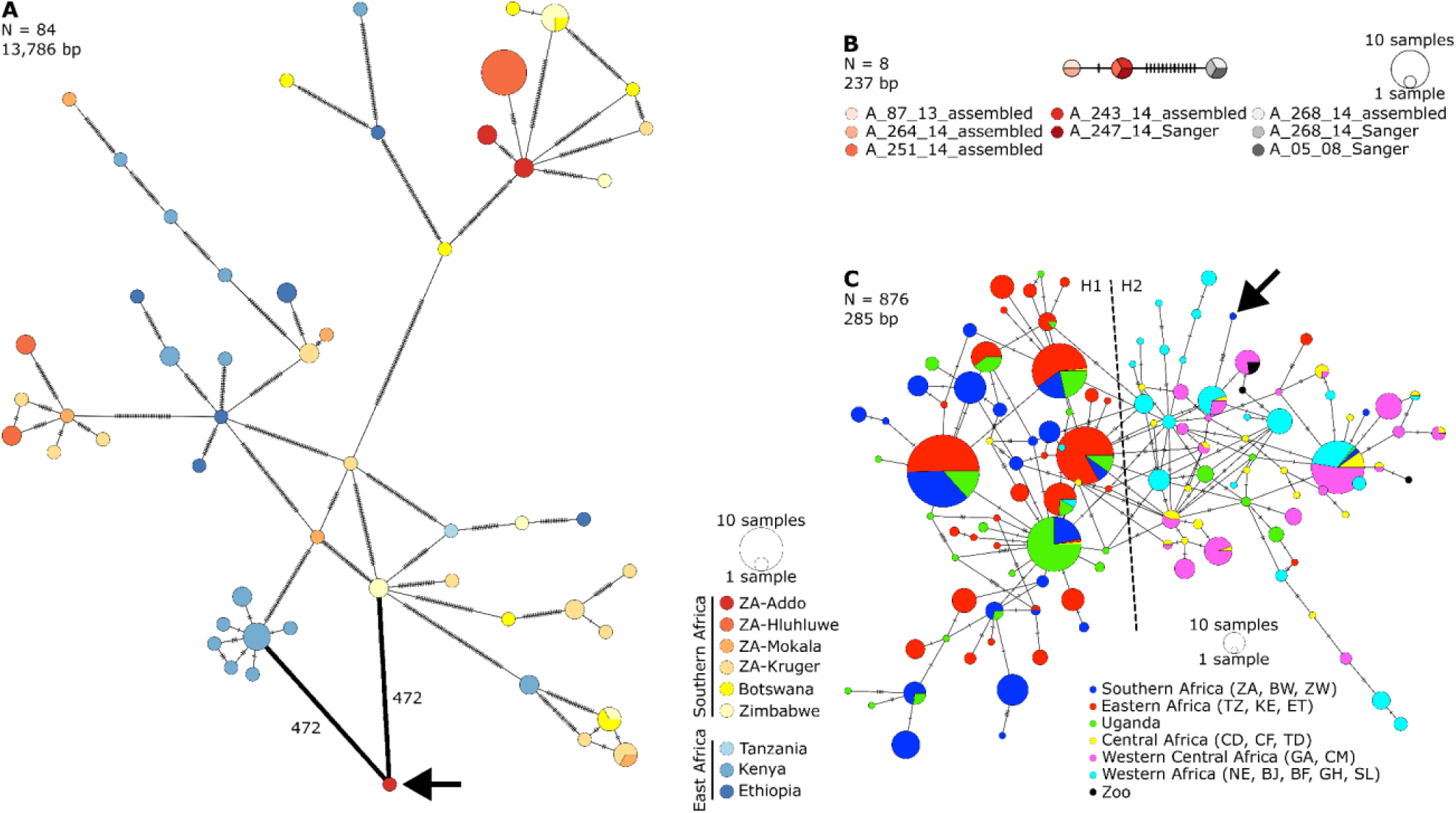
Minimum spanning haplotype networks of: **A** the 13,786 bp alignment of 13 protein-coding genes, rRNA genes, and CR from 44 publicly available and 40 newly generated Cape buffalo mitogenomes, **B** the 237 bp alignment of cytochrome *b* of Addo samples from both assembled mitogenomes and those generated via Sanger sequencing, and **C** the 285 bp alignment of publicly available CR sequences of all African buffalo subspecies and from the 40 newly generated mitogenomes in this study. The colours in **C** follow those from Smitz et al. (2013) to facilitate comparison with their Figure 1. Each circle represents a unique haplotype; the sizes of the circles represent the frequencies as per the scales shown. Cross hatches on links connecting haplotypes each represents one substitution. The arrows in **A** and **C** indicate the divergent mitogenome from Addo (sample A_268_14). The numbers in **A** show the number of mutational steps between the divergent mitogenome and the nearest haplotypes. ISO 3166 two-letter country codes used in **A** and **C** refer as follows: ZA – South Africa, BW – Botswana, ZW – Zimbabwe, TZ – Tanzania, KE – Kenya, ET – Ethiopia, CD – The Democratic Republic of the Congo, CF – Central African Republic, TD – Chad, GA – Gabon, CM – Cameroon, NE – Niger, BJ – Benin, BF – Burkina Faso, GH – Ghana, SL – Sierra Leone. “Zoo” refers to samples from Antwerpen Zoo (Belgium, n = 2), Berlin Zoo (Germany, n = 1), Dresden Zoo (Germany, n = 1), and Safari Park Beekse Bergen (Netherlands, n = 1) (van Hooft et al., 2002). All networks were generated in PopArt and further processed in InkScape.

### A highly divergent mitogenome in Addo Elephant NP

The divergent mitogenome from an Addo Elephant NP buffalo encoded full length proteins with no internal stop codons, thus was likely not an assembly error or a NuMT. This mitogenome did not cluster with the others from Addo Elephant NP in the haplotype network (Figure 1A) or the phylogenetic tree (Figure S1). The closest haplotypes were the most common haplotype in Kenya and another from Zimbabwe, although there were 472 substitutions between each of these and the divergent mitogenome. To further validate the authenticity of the divergent mitogenome, we performed Sanger sequencing of a fragment of the cytochrome *b* gene of this sample (A_268_14, male), as well as its two closest relatives, A_05_08 (male, Wang pairwise relatedness estimate (*r*) = 0.45) and A_247_14 (female, *r* = 0.31), and constructed a haplotype network with the cytochrome *b* genes of other assembled Addo mitogenomes (Figure 1B). We found that the Sanger-generated sequence of A_268_14 was identical to that of the assembled sequence, indicating again that it was unlikely to be an assembly error or NuMT. Furthermore, its closest relative (A_05_08) had an identical Sanger-generated sequence and thus shares this divergent mitogenome with sample A_268_14. Two other haplotypes were identified, separated from the divergent haplotype by 13 and 14 mutations across 237 bp, which were shared among the five other Addo samples, including A_247_14, the next-closest relative of A_268_14 after A_05_08 (Figure 1B).

The divergent mitogenome might have a relatively high frequency in the Addo population, as it was present in one of five (20%) unrelated individuals selected for whole genome sequencing in de Jager et al. (2021). This frequency increases to two in seven (28.5%) if the Sanger sequencing results are included, though this sampling was biased towards close relatives of A_268_14. Nevertheless, the sample size was relatively low compared to the census size of Addo (∼800 buffalo) and thus likely does not adequately represent the frequency of the divergent haplotype in the population. However, it is evidently at a high enough frequency to be detected from just five samples. The Addo population experienced a relatively severe bottleneck in the late 1800s and early 1900s, and remained at low numbers (<250) throughout much of the 1900s, with the lowest known census size being 52 individuals in 1985 (*Pers. Comm.* D. Zimmerman 2015; (de Jager et al., 2021)), resulting in the loss of genetic diversity through drift and inbreeding (de Jager et al., 2021; de Jager et al., 2020; O’Ryan et al., 1998; Quinn et al., 2023). Despite this, the divergent mitochondrial lineage was maintained in the population, pointing towards a high pre-bottleneck frequency. Addo Elephant NP buffalo have been used to seed and supplement other buffalo populations in southern Africa and have been sold to private wildlife ranchers for many years, raising the possibility that the divergent lineage may also now be present in those populations and ranches. Future studies could robustly estimate the frequency of this haplotype in the Addo population through more extensive sampling and investigate whether there are any potential fitness effects for individuals that carry this divergent mitochondrial lineage.

To determine where the divergent mitogenome clusters within the diversity of the entire species, we constructed a minimum spanning haplotype network of 285 bp of the CR sequence from all African buffalo subspecies (Figure 1C). We were able to reconstruct lineages H1 and H2 as identified in Smitz et al. (2013), where H1 is the South-Eastern lineage (mainly Cape buffalo, *S. c. caffer*) and H2 is the West-Central lineage (predominantly containing the other three subspecies, *S. c. nanus*, *S. c. aequinoctialis*, *S. c. brachyceros*). The lineages are not monophyletic, and each contains some discordant haplotypes (Figure 1C, also see Figure 1 in Smitz et al. (2013)). The CR haplotype of the divergent Addo sequence (of *S. c. caffer* origin) grouped with the H2 (West-Central) lineage and not the H1 (South-Eastern) lineage like most other *S. c. caffer* sequences (Figure 1C). It was at a tip of the network, separated by five substitutions from a relatively common haplotype from Western Africa that was also present in West Central, and Central Africa (Figure 1C). Its location in the H2 lineage, and the number of substitutions separating it from its nearest haplotype, is not especially unique in this network. There are several haplotype-lineage “mismatches” and other haplotypes that are separated by four or six substitutions, the latter always involving haplotypes from Uganda (Figure 1C). This is interesting, because it highlights the limited power of using only a portion of the CR, as high diversity in this region might be expected and the highly divergent nature of the rest of the mitogenome of lineages such as the one in Addo Elephant NP may go undetected or under-investigated. It also raises the question of whether the other relatively diverged haplotypes in Figure 1C (from Uganda) are as highly divergent across the rest of the mitogenome as the Addo mitogenome is, or whether the variation is localised to the CR, as is the case with the Ethiopian mitogenome (ID: 9083) in Heller et al. (2012) with a 79 bp deletion in the CR, but it still falls within the Cape buffalo diversity (Figure S1).

Finally, using a fossil-calibrated phylogeny, we found that the divergent Addo mitogenome formed a sister lineage to other Cape buffalo, with a divergence date of approximately 2.51 million years ago (Mya) (95% highest posterior density: 1.89-3.29 Mya) (Figure 2). This estimate overlaps with the divergence time of both species-level divergence [bison (*Bison bison*) and yak (*Bos grunniens*)], and subspecies-level divergence (water buffalo subspecies: *Bubalus bubalis bubalis* and *B. b. carabanesis*) within Bovini (Figure 2), an observation which also holds within the Bovidae family (Bibi, 2013). All divergence dates of known species-level relationships estimated here (Figure 2) correspond to accepted, published estimates based on mitogenomes (Bibi, 2013) and whole nuclear genomes (Chen et al., 2019), indicating that the divergence date of the Addo mitogenome of interest is most likely accurate.

**Figure 2.**
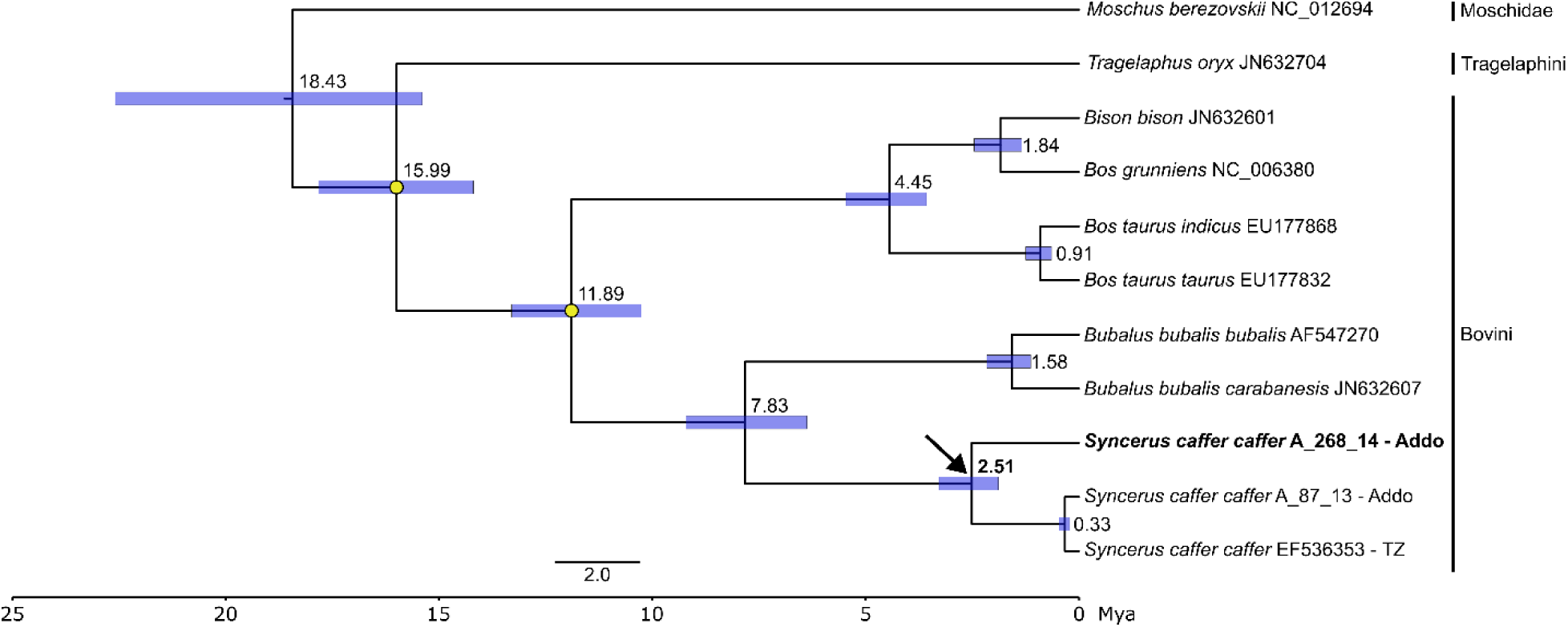
Fossil-calibrated, Bayesian phylogenetic tree of mitogenomes estimated in BEAST. Yellow circles show the fossil calibration points used. Numbers at the nodes show the estimated age of the node in millions of years. Node bars represent the 95% highest posterior density of the node age. All nodes have a posterior support of 1. The divergent Addo mitogenome is highlighted in bold text, with the arrow indicating the node where it diverged from other Cape buffalo mitogenomes. TZ – Tanzania, Mya – million years ago.

### Source of the divergent Addo mitogenome

Given the high level of divergence (3.5% and 2.51 Mya) of the divergent Addo mitogenome from other Cape buffalo mitogenomes, we hypothesise that the most likely source is through introgression with a closely related species or subspecies. The alternative hypothesis of a population-specific divergence event, likely would have resulted in a divergence time of younger than 449,000 years – the current maximum estimated divergence date of extant African buffalo lineages (Smitz et al., 2013). In the phylogenetic analysis, the divergent mitogenome unequivocally grouped with Cape buffalo (node posterior support = 1), which excludes water buffalo (*Bubalus bubalis*) and cattle (*Bos taurus*) as potential sources. Importantly, we note that there have been no recorded human-mediated introductions of African buffalo into the Addo population since it was proclaimed as a protected area in 1931 (*Pers. Comm.* D. Zimmerman 2015; (de Jager et al., 2021)), and it therefore represents the long-term, naturally occurring population and lineages of Cape buffalo in that region of South Africa.

A potential source of the divergent mitogenome could be introgression from other extant African buffalo subspecies in eastern Africa before *S. c. caffer* dispersed to southern Africa. Unfortunately, no mitogenomes or mitochondrial gene sequences from any of the other three subspecies were publicly available to test this possibility, and while the CR haplotype network may lend some support to this, it is based on very limited data and thus may be misleading. Furthermore, the recent split between the two African buffalo lineages (H1: *S. c. caffer* and H2: *S. c. nanus*, *S. c. aequinoctialis*, *S. c. brachyceros*) of between 145,000 and 449,000 years ago (Smitz et al., 2013) indicates that another extant subspecies is unlikely to be the source of the divergent mitogenome. There are no other extant bovines that occur in Africa or around Addo Elephant NP that might be the source of this mitogenome through introgression.

The present-day southern African buffalo populations are likely the result of a population expansion event from eastern to southern Africa between 80,000 and 50,000 years ago (Heller et al., 2012; Smitz et al., 2013; van Hooft et al., 2002). However, there is radiometric evidence from Kathu Pan in the Northern Cape Province of South Africa that fossils from this site are around 542,000 years old, which includes *Syncerus caffer*, *S*. *antiquus* (the extinct long-horned buffalo), and other undetermined buffalo specimens (Porat et al., 2010). Consequently, Smitz et al. (2013) proposed the possibility that dispersal into southern Africa occurred multiple times throughout the Mid-to Late Pleistocene (∼1.25-0.0117 Mya). It may thus be that the divergent mitogenome could be from a remnant population of *S*. *caffer* from one of these previous dispersal events that survived around present-day Addo and was assimilated into the contemporary population during subsequent dispersal events. However, the young origin of the extant buffalo subspecies (see above) and of *S*. *caffer* as a species, which first appears in the fossil record ∼1 Mya at Nariokotome in the Turkana Basin in Kenya (Faith, Rowan, & Du, 2019; Fortelius et al., 2016), would preclude the above scenario as too recent to be the source of the divergent mitogenome.

Thus, the most likely origin of the divergent mitochondrial lineage appears to be an ancient introgression event with an extinct buffalo species. Two extinct *Syncerus* species have been described from the fossil record, *S. acoelotus* and *S. antiquus*, while many fossil specimens are only assigned to the genus level. The genus first appears in the fossil record during Late Pliocene (∼3.6-2.58 Mya) in eastern Africa, with specimens assigned to *Syncerus* cf. being dated to ∼2.8 Mya at the Shungura Formation in the Omo Valley in Ethiopia (Faith et al., 2019). *Syncerus acoelotus* occurs in the fossil record from ∼2.7 Mya (Taung, South Africa (McKee, 1993)) to ∼0.7 Mya (Olduvai Gorge, Tanzania (Gentry & Gentry, 1978)). *Syncerus antiquus* appears at around the same time as *S. caffer* at ∼1 Mya in the Bouri Formation at the Daka site in Ethiopia (Faith et al., 2019; Gilbert & Asfaw, 2008) and is suggested to have gone extinct at the end of the Pleistocene (∼0.0117 Mya (Klein, 1984; Klein, 1994)) south of the Sahara Desert and ∼0.004 Mya in North Africa (Gautier & Muzzolini, 1991). Thus, both extinct *Syncerus* species overlapped in time and space with the extant *S. caffer*, providing opportunity for introgressive events to occur.

This hybridisation and introgression could have occurred in eastern Africa between either of the extinct species and the ancestral *S. caffer*, before the emergence of the extant subspecies, and this variation was retained until the present day and detected in the Addo Elephant NP population. A similar scenario has been described in the spiral-horned antelope (Rakotoarivelo, O’Donoghue, Bruford, & Moodley, 2019). Similarly, divergent lineages may exist in other extant populations, such as in Uganda where equally divergent CR sequences were observed (Figure 1C), but from which mitochondrial gene sequences or whole mitogenomes are unavailable at present.

Alternatively, though not necessarily to the exclusion of the previous scenario, as *S. caffer* (or *S. c. caffer*) dispersed into southern Africa it may have come across and interbred with either *S. acoelotus* or *S. antiquus* populations already present in the region, with the mitochondrial lineages from those introgressive events being retained in southern African populations. As indicated above, *Syncerus* has a long history in southern Africa (at least since 2.7 Mya) and occurs at various sites in the region throughout the Pleistocene (Avery, 2019). Additionally, Cape (*S. c. caffer*) and long-horned (*S. antiquus*) buffalo co-occurred in the Cape region of South Africa, as evidenced from palaeontological sites in South Africa, near Addo Elephant NP, namely Klasies River Mouth Caves, Nelson Bay Cave, and Byneskranskop Cave 1 (Klein, 1994), providing opportunity for hybridisation and introgression in the region of the present-day Addo Elephant NP population. It should be noted that no intermediate (hybrid) forms have been found in the fossil record (Klein, 1994). However, this may be due to the close resemblance of their postcranial anatomy (e.g., Peters, Gautier, Brink, & Haenen (1994) classified the long-horned buffalo as a subspecies of *S. caffer*: *S. c. antiquus*) making it difficult to identify hybrids, especially if back-crossing occurs with either parent, or because fossilisation and hybridisation are both rare events, making the preservation and discovery of hybrid fossils likely an extremely rare event.

While the fossil record indicates that there potentially was ample opportunity for *S. c. caffer* and *S. antiquus* to interbreed, it does not align with the divergence date obtained with the mitogenome (2.51 Mya, Figure 2), as both species only appear ∼1 Mya. Consequently, the dated fossil record rather supports *S. acoelotus*, appearing ∼2.7 Mya, as the source of the mitogenome through introgression with *S. caffer* or *S. c. caffer*. Of course, the fossil record does not provide a complete species history and both *S. caffer* and *S. antiquus* may be older than is currently accepted, or the source may be an undescribed or undetected extinct *Syncerus* species. Finally, there is no clear reason, to our knowledge, why the introgressive hybridisation process (if that is indeed what has occurred) should have been localised to the southern tip of Africa, as the three *Syncerus* species discussed co-occurred in space and time across a much larger part of Africa (Faith, 2014). It could be that it has simply not been detected yet, due to larger population sizes in other regions compared to Addo Elephant NP, which would require a more intensive sampling effort to detect a rare lineage, whereas in Addo the lineage appears to be at a relatively high frequency.

Several opportunities exist to gain some clarity regarding the source of the divergent mitogenome in the Addo Elephant NP population. First, whole mitogenome sequences from all extant African buffalo subspecies, and particularly from Ugandan populations, would provide better context to determine whether the divergent mitogenome falls within or outside the diversity of *S. caffer*. Second, mitochondrial DNA sequences from the extinct *S. antiquus* would allow this species to be confirmed or excluded as the source of the divergent mitogenome, and if it is excluded, would potentially provide support that *S. acoelotus* was the source. With the techniques in the fields of ancient DNA and paleogenomics constantly improving, it is now possible to obtain these ancient genetic data for *S. antiquus*, as shown by Hempel et al. (2022) who recovered the nuclear genome of the extinct blue antelope (*Hippotragus leucophaeus*) from a specimen dating to 9,800-9,300 years old from Nelson Bay Cave in South Africa. Several *S. antiquus* specimens are available from the end of the Pleistocene (∼23,000-12,000 years ago) from Nelson Bay Cave (Loftus, Sealy, & Lee-Thorp, 2016; Klein, 1972) and Boomplaas Cave (Faith, 2013) in South Africa, which could be a good starting point for to attempt to obtain *S. antiquus* ancient DNA.

### Conservation consequences for the Addo Elephant NP buffalo population

Regardless of the source of the divergent mitogenome, the fact that it is found in the present-day Addo buffalo population further increases the conservation value of this population and indicates that its genetic management requires careful consideration. This population already has high conservation value in South Africa for several reasons: It is one of three remnant populations that survived the hunting onslaught and rinderpest epidemic in the 1800s and 1900s (the others being Kruger National Park and Hluhluwe-iMfolozi Park); it is the only naturally occurring disease-free population in the country (another disease-free population of Kruger National Park buffalo was established in Mokala National Park and Graspan); and finally, Addo buffalo are used to seed or supplement populations throughout southern Africa and are sold to private wildlife ranchers (de Jager et al., 2020; Laubscher & Hoffman, 2012). However, it has long been suspected and repeatedly shown that this population has low genetic diversity due to the previously-mentioned population bottleneck and subsequent isolation, and that it is threatened by inbreeding (de Jager et al., 2021; de Jager et al., 2020; O’Ryan et al., 1998; Quinn et al., 2023).

To prevent further inbreeding and inbreeding depression (the manifestation of inbreeding in the biology of the population or individuals, such as reduced population growth, increased susceptibility to disease, low sperm quality, low body condition, etc.), O’Ryan et al. (1998) calculated that one breeding bull (male) of Kruger stock should be introduced to the Addo population every generation (∼7.5 years), which can now be done via the disease-free population in Mokala NP (de Jager et al., 2020). This frequency of introduction was proposed to prevent genetic swamping. Wisely, it was also suggested that any introduced individuals be males (O’Ryan et al., 1998). If this guidance is followed, any unique mitochondrial diversity, such as the highly divergent lineage identified here, should be retained in the Addo population (if any female buffalo carry this lineage), because mitochondrial DNA is inherited only from mother to offspring.

## Conclusion

The findings in this study highlight the importance of conserving distinct genetic units within a species or subspecies and the value of a cautionary approach to genetic management of populations. Previous genetic studies based on nuclear DNA found that while Addo, Hluhluwe-iMfolozi and Kruger buffalo populations represent separate genetic clusters, it did not seem that there was much unique diversity within Addo, and that it contained a subset of the genetic variation found in Kruger (de Jager et al., 2021; de Jager et al., 2020; O’Ryan et al., 1998; Quinn et al., 2023). This is true to a certain extent when looking at nuclear DNA, but this study has shown that all three South African populations contain unique mitochondrial lineages, and that the Addo population contains this highly divergent, cryptic mitochondrial lineage that has not yet been detected in any other Cape or African buffalo populations. Whether this mitochondrial lineage confers any fitness or adaptive advantage to Addo buffalo is not yet known, but it has persisted in the population for potentially thousands of years alongside other, more common, Cape buffalo lineages, and this further increases the conservation value of this vulnerable population. If the Addo population were allowed to go extinct, or swamped with Kruger buffalo, this lineage would probably have been lost. Thus, an intermediate position, where genetic clusters are preserved as far as possible, but inbreeding depression is prevented, emerges as a sensible approach to genetic management of these populations.

## Supplementary material

**Supplementary Table S1.** Metadata of mitogenomes used in this study.

**Supplementary Table S2.** Metadata of control region sequences used in this study.

**Supplementary Table S3.** Assembly statistics of the mitogenomes assembled in this study with NOVOPlasty v4.0 from the reads of whole nuclear genomes sequenced by de Jager et al. (2021).

**Supplementary Figure S1** Bayesian phylogenetic tree of Cape buffalo whole mitogenomes (16,436 bp alignment) constructed in MrBayes. The cattle (Bos taurus) and water buffalo (Bubalus bubalis) reference mitogenomes were used as outgroups. Branches are coloured by posterior probability, as indicated by the colour scale bar. Posterior probabilities for nodes are only shown if less than one, with the exception of the node supporting the divergent Addo mitogenome (indicated by the arrow). Sample names are coloured by country of origin, and by protected area of origin in the case of South African samples, as in Figure 1A. Sample names are followed by the ISO 3166 two-letter code of the country of origin, which refer as follows: ZA – South Africa, BW – Botswana, ZW – Zimbabwe, TZ – Tanzania, KE – Kenya, ET – Ethiopia. Those samples from South Africa (ZA) also followed by the protected area of origin.

## Acknowledgements

The Centre for High Performance Computing (CHPC), Cape Town, South Africa, is hereby acknowledged for providing the computing resources used to conduct the analyses in this study, as well as the Mjolnir computing cluster at the Globe Institute, University of Copenhagen. The DNA Sequencing Facility of the University of Pretoria is acknowledged. We further acknowledge the support of the DSI-NRF Centre of Excellence for Biomedical Tuberculosis Research, South African Medical Research Council Centre for Tuberculosis Research, Division of Molecular Biology and Human Genetics, Faculty of Medicine and Health Sciences, Stellenbosch University, Cape Town, South Africa. DdJ acknowledges postdoctoral research funding from the University of Pretoria and the National Research Foundation (grant ID: 111962) and funding from the IRT Genomics, University of Pretoria. Opinions, findings and conclusions, or recommendations expressed herein are those of the researchers and do not necessarily reflect those of the funding bodies. DdJ thanks Louis du Plessis and Tyler Faith for valuable insight into the manuscript after posting on the bioRxiv preprint server.

