## Supplementary figures and images for "A highly divergent mitochondrial genome in extant Cape buffalo from Addo Elephant National Park, South Africa"

### Figure S1

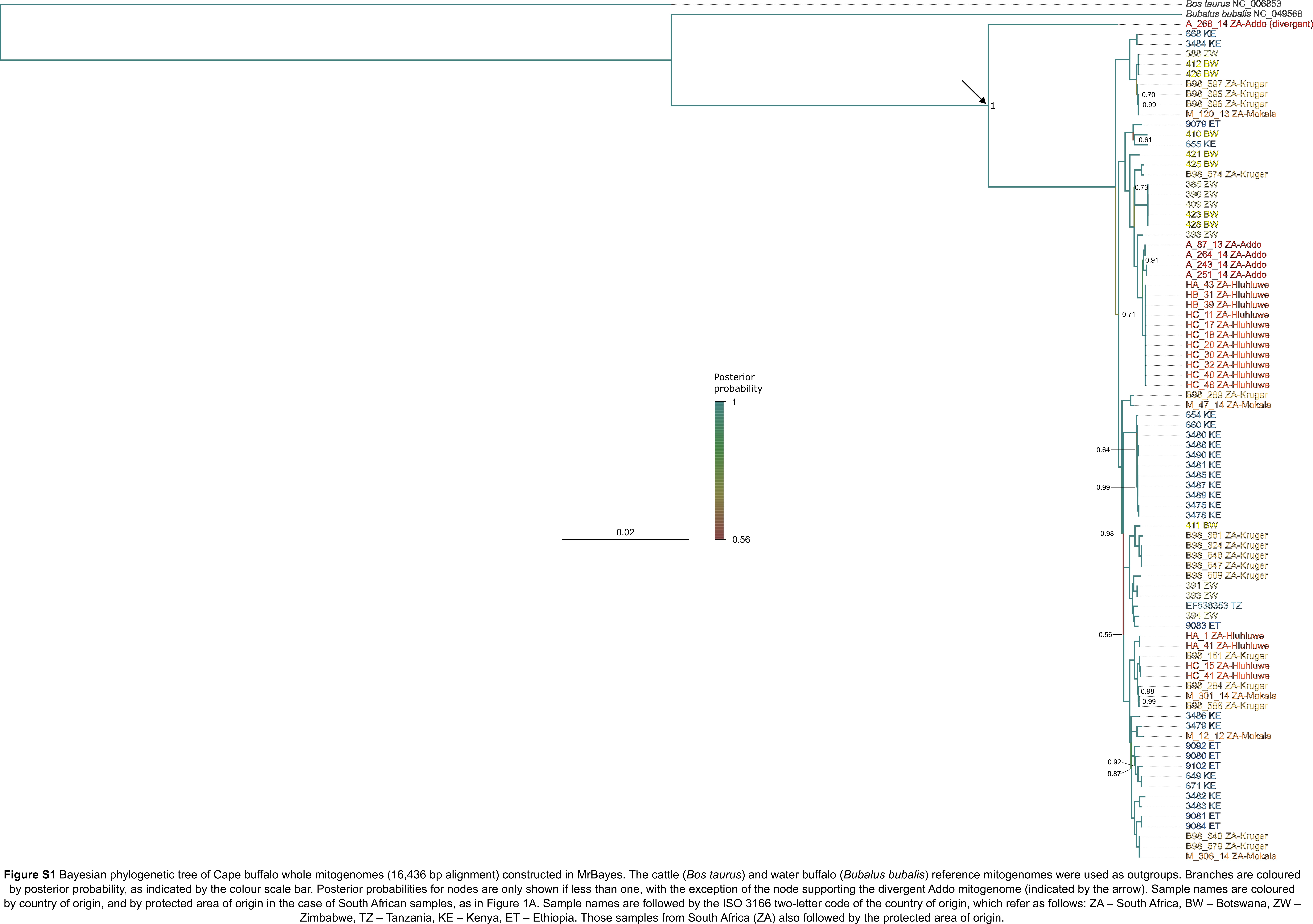
